# Characterising the association between posterior parietal metabolite levels and cortical macrostructure in a cohort spanning childhood to adulthood

**DOI:** 10.1101/2025.05.20.655054

**Authors:** Alice R Thomson, Duanghathai Pasanta, Richard A E Edden, Tomoki Arichi, Xiaoqian Chai, Nicolaas A Puts

## Abstract

Postnatal brain development is characterised by dynamic macrostructural changes, including cortical thinning and cortical flattening during childhood and adolescence. These macro-structural changes are parallel with developmental changes in brain neurochemistry, probed in the human brain using Magnetic Resonance Spectroscopy (MRS). This includes neurotransmitters such glutamate and gamma-aminobutyric acid (GABA), as well as building blocks of neuronal and associated tissue such as N-acetyl aspartate (NAA), and those involved in metabolism such as creatine (Cr). While previous research has linked MRS-measured neuro-metabolite levels to bulk tissue composition (e.g., gray matter, white matter, and cerebrospinal fluid), the relationship between neurochemistry and more granular macrostructural metrics, such as cortical thickness, area, volume, and local gyrification, remains unexplored. This study investigates the association between MRS-measured neuro-metabolite levels in the posterior parietal cortex (PPC) and PPC-voxel cortical macrostructural metrics in a developmental cohort of 86 individuals aged 5–35 years. We also examine whether PPC metabolite concentrations associate with whole-brain structural metrics to determine whether associations are region-specific or more broadly generalisable. Our findings reveal significant positive associations between PPC cortical thickness, volume, local gyrification index (LGI) and Glx (glutamate + glutamine) levels, likely because differences in cortical *microstructure*, including dendritic arbour complexity, contributes to variation in both cortical macrostructure and Glx activity across development. Additionally, PPC Glx:GABA+ ratio negatively associated with subcortical gray matter volume, while PPC total NAA positively associated with cerebral white matter volume, suggesting a link between regional neurochemistry and broader brain structure. These results highlight the importance of accounting for macrostructural and broader brain structural characteristics when interpreting the neuroanatomical correlates of MRS-measured metabolites, beyond controlling for bulk tissue composition. This approach is particularly crucial when comparing neuro-metabolite levels across groups with known structural differences, such as developmental cohorts or individuals with neurodevelopmental conditions.

**Key points:**

- Posterior parietal cortex (PPC) Glx levels are positively associated with PPC cortical thickness and local gyrification index, likely because differences in cortical microstructure, including dendritic arbour complexity, contributes to both cortical macrostructure and neuro-metabolic traits across development.
- The PPC Glx:GABA+ ratio is negatively associated with subcortical gray matter volume, while PPC total NAA is positively associated with cerebral white matter volume, suggesting a link between regional neurochemistry and broader brain structure.
- These findings emphasise the importance of considering more detailed macrostructural characteristics, as well as bulk tissue composition (white matter, gray matter, cerebral spinal fluid), when interpreting MRS-measured metabolite differences.

## Introduction

Postnatal brain development is characterised by dynamic changes in brain macro-structure from childhood to adulthood, including cortical thinning (Baik et al., 2023; Ducharme et al., 2016; Frangou et al., 2022; Koolschijn & Crone, 2013; Muftuler et al., 2011) and decreases in cortical folding (Alemán-Gómez et al., 2013; Ducharme et al., 2016; Klein et al., 2014; White et al., 2010). Cortical thinning during childhood and adolescence is thought to reflect developmental remodelling of laminar structural complexity driven by dendritic arbour pruning (la Fougère et al., 2011; Petanjek, Judaš, Šimic, et al., 2011; Sowell et al., 2004; Vidal-Pineiro et al., 2020). Synaptic connections in the cortex are initially formed in excess, peaking in early childhood (2-4 years of age; Huttenlocher, 1979), before weaker connections are subsequently eliminated from childhood to early adulthood (Faust et al., 2021; Huttenlocher, 1979; Huttenlocher & Dabholkar, 1997; Huttenlocher & de Courten, 1987; Selemon, 2013). This achieves highly efficient, adaptive and precise connectivity, as the fewer remaining synapses are strengthened and maintained (Bourgeois et al., 1989, 1994; Geschwind, 2011; Huttenlocher & Dabholkar, 1997; Huttenlocher & de Courten, 1987, 1987; Petanjek, Judaš, Šimic, et al., 2011; Selemon, 2013; Zecevic et al., 1989). This process of structural refinement in the cortex is closely linked to cognitive development (Fleming & McDermott, 2024). Similarly, gyrification or cortical folding is a developmental process whereby the cortical surface is folded, increasing cortical surface area within a fixed volume to support efficient structural connectivity (White et al., 2010), and making functionally distinct brain areas spatially isolated (Jiang et al., 2021). Cortical folding predominantly occurs pre- and early post-natally (White et al., 2010). Gradual decreases in local gyrification index (LGI), a measure of cortical folding (lower LGI indicating a lesser degree of cortical folding), during childhood and adolescence have however been linked to changes in dendritic morphology and synaptic connectivity also, which alters cortical structural connectivity and so folding forces, leading to decreased sulcal depth and flattening of the cortex (Alemán-Gómez et al., 2013; White et al., 2010).

Such developmental changes in brain macro-architecture are concurrent with developmental changes in brain neurochemistry, probed in the human brain using Magnetic Resonance Spectroscopy (MRS). This includes neurotransmitters such as gamma-aminobutyric acid (GABA) and glutamate, as well as building blocks of neuronal and associated tissue such as N-acetyl aspartate (NAA), choline (Cho), and those involved in metabolism such as creatine (Cr) and myo-inositol (mI, Rae, 2014). Recent studies suggest that MRS metabolites undergo developmental shifts that parallel changes in cortical macrostructure across childhood and adolescence (Perdue et al., 2023; Reyngoudt et al., 2012; Thomson, Hwa, et al., 2024), while strong associations between cortical thickness and genes related to neuronal metabolism and function further support a potential link between these macrostructural and neurochemical changes (Z. Zhou et al., 2023).

Despite these findings, the association between changes in neurochemistry and finer-grained structural metrics, such as cortical thickness, area, volume and cortical folding (estimated as LGI), has yet to be explored in a single typical cohort. Prior studies have primarily established links between neuro-metabolite concentrations and bulk voxel tissue composition (gross gray matter, white matter and CSF), limiting the ability to draw detailed conclusions about the structural basis of metabolite differences. Macrostructural metrics, such as cortical thickness and LGI, provide more detailed insights into cortical architecture, and developmental changes in these measures are likely to interact with underlying biochemical processes, shaping regional but also *global* brain development. Although brain regions mature at different rates, they are highly interconnected and influence one another’s development (Antón-Bolaños et al., 2018; Casey et al., 2008; Gogtay et al., 2004; Kingsbury et al., 2002; L. Zhou et al., 2010). For instance, subcortical thalamic afferents developmentally modulate cortical neuron activity, essential for typical cortical structural development, including dendritic morphology, synaptic density and cortical topography (Antón-Bolaños et al., 2018; Kingsbury et al., 2002; L. Zhou et al., 2010). Thus, regional brain neurochemistry may interact with wider brain structural development, a connection that also remains underexplored.

To address this gap, this study is the first to investigate the associations between MRS-measured neuro-metabolite levels in the posterior parietal cortex (PPC), which we previously characterised (Thomson, Hwa, et al., 2024), and macro-structural metrics derived from the same PPC MRS voxel in a developmental cohort spanning 5 to 35 years of age. To determine whether the identified associations are region-specific or globally relevant, we also investigate interactions between PPC metabolite concentrations and whole brain structural metrics including cerebral white matter volume and subcortical gray matter volume. The PPC voxel was selected due to its more protracted development (Hill et al., 2010; Kaas & Stepniewska, 2016), enabling us to associate developmental changes in macrostructure and neurochemistry from mid-childhood through to early adulthood. Understanding the neuroanatomical correlates of changes in regional brain chemistry across this period is not only valuable for a more detailed characterisation of the typical brain development, but also provides a more comprehensive framework for understanding the biological underpinnings of neurodevelopmental disorders where neurochemistry *and* macrostructure have been shown to be disrupted (Ecker et al., 2016; Hyde et al., 2010; Kohli et al., 2019; Puts et al., 2017; Thomson, Pasanta, et al., 2024) and may be associated.

## Methods

### Participants

117 participants underwent in vivo MRI imaging, overseen through local Kennedy Krieger Institute and Johns Hopkins University IRB procedures. Children assented, and their parents consented, to participation. All participants were native English speakers, right-handed, had normal or corrected-to-normal vision, with no history of psychiatric, neurological, or developmental disorders. All participants had IQ > 85. Participant demographic data were collected at the time of scan. The study received approval through the local IRB procedures at the Kennedy Krieger Institute and Johns Hopkins University, with all participants providing informed consent prior to their involvement. 86 of these participants, with high-quality MRS and structural data (see details below), were included in this analysis, aged between 5 and 35 years of age (Table 1). The range of participant IQ scores was 85–138 (median IQ = 117). Data quality was assessed both across and within age groups (children, adolescents, and adults) to examine potential quality bias associated with age (see Table 1).

**Table 1.** Sample demographics by age group. Note only participants with high-quality MRS and structural MRI data were included.

|  | *N*<br>*(females/males)* | *Median age*<br>*(IQR)* | *median IQ*<br>*(IQR)* |
| --- | --- | --- | --- |
| *Child ($5 \geq \text{age} < 12$)* | 31 (15/16) | 10.135 (2.58) | 115 (17.25) |
| *Adolescent ($12 \geq \text{age} < 18$)* | 22 (10/12) | 14.41 (2.25) | 113 (15.00) |
| *Adult ($\text{age} \geq 18$)* | 33 (15/18) | 21.67 (7.67) | 120.00 (10.00) |
| *All participants* | 86 (40/46) | 14.71 (9.52) | 117.00 (16.25) |

### MRI

#### Structural data

MRI imaging was performed on a Philips 3 Tesla Achieva scanner (Best, NL) at the F.M. Kirby Research Centre for Functional Brain Imaging at the Kennedy Krieger institute in Baltimore, USA with a 32-channel receive head coil. High-resolution (1 mm^3^ isotropic) T_1_-weighted magnetisation-prepared rapid acquisition gradient-echo sequence (MPRAGE) anatomical images were acquired (slice thickness; 0.83 mm; in-plane resolution, 1×0.83 mm; TR 7 ms; TE 3.2 ms), reconstructed and visually inspected for artifacts (by AT).

#### MRS data

MRS data reported here have been reported previously (Thomson, Hwa, et al., 2024) per reporting standards (Lin et al., 2021), and were acquired from a 27 ml voxel (3 × 3 × 3 cm^3^) placed over the posterior parietal cortex (PPC) and centred on the midline (Figure 2). MRS was performed using GABA+-selective MEshcher-Garwood Point RESolved Spectroscopy (MEGA-PRESS; (Mescher et al., 1998); 320 transients (160 ON and 160 OFF), 2048 data points, TE/TR 68/2000 ms with editing pulses placed at 1.9 ppm in the edit-ON acquisitions and 7.46 ppm in the edit-OFF acquisitions, and using VAPOR water suppression. An interleaved unsuppressed water reference with the same parameters for water supressed scans was used (16 averages) to mitigate scanner drift and for subsequent eddy-current and phase corrections, and metabolite quantification (Edden et al., 2016). Thirty-one MRS datasets were excluded due to poor data quality (for details see Thomson, Hwa, et al., 2024). As such, our MRS and structural data analysis uses data from 86 participants. Demographics for this analysis cohort are shown in Table 1.

### Structural data processing

T_1_-weighted images were motion corrected, transformed and intensity corrected in FreeSurfer (version 7.4.1). Images were segmented (into GM, WM, CSF volumes) and the cortical surface reconstructed (Dale et al., 1999; Fischl et al., 1999; Fischl & Dale, 2000). The FreeSurfer commands *recon-all* and *mris_compute_lgi* were used to calculate local gyrification index (LGI) for each point on the cortical surface (typically scoring between 1 and 5). First, a smooth outer surface mask, enveloping the pial surface, is created for each participant. The smooth outer surface mask and the pial surface are segmented into smaller, identical circular regions from which the local gyrification is calculated for each vertex, this being the ratio of the area of the pial surface at the outer smooth mask to the smooth outer surface mask (Schaer et al., 2008). Whole-brain measures including subcortical gray matter volume (e.g. sum of the gray matter voxel volumes, in mm^3^), cerebral white matter volume (in mm^3^), cortical volume (in mm^3^), and estimated intracranial volume (eTIV, in mm^3^), were extracted from the results of the FreeSurfer procedures for each participant.

#### Extracting voxel surface metrics

The Freesurfer command *mri_vol2surf* was used to project the PPC MRS voxel mask onto the cortical surface (pial). This was performed individually for each participant as voxel placement varied based on individual brain anatomy. As the MRS PPC voxel was centred to the midline, the voxel volume was projected into the medial parts of both the right and left hemisphere, creating two voxel masks (see Figure 1). The Freesurfer *MRI_segstats* command was then used to extract quantitative measures from the 2D voxel masks projected onto the inflated right and left hemisphere cortical surface. Measures extracted included mean cortical thickness, which is the shortest distance between the pial and white matter surface (mm), cortical volume (mm^3^) and voxel LGI (Schaer et al., 2008). Per PPC voxel structural metric, mean values were calculated from right and light hemisphere estimates and used for all analysis described below.

**Figure 1.**
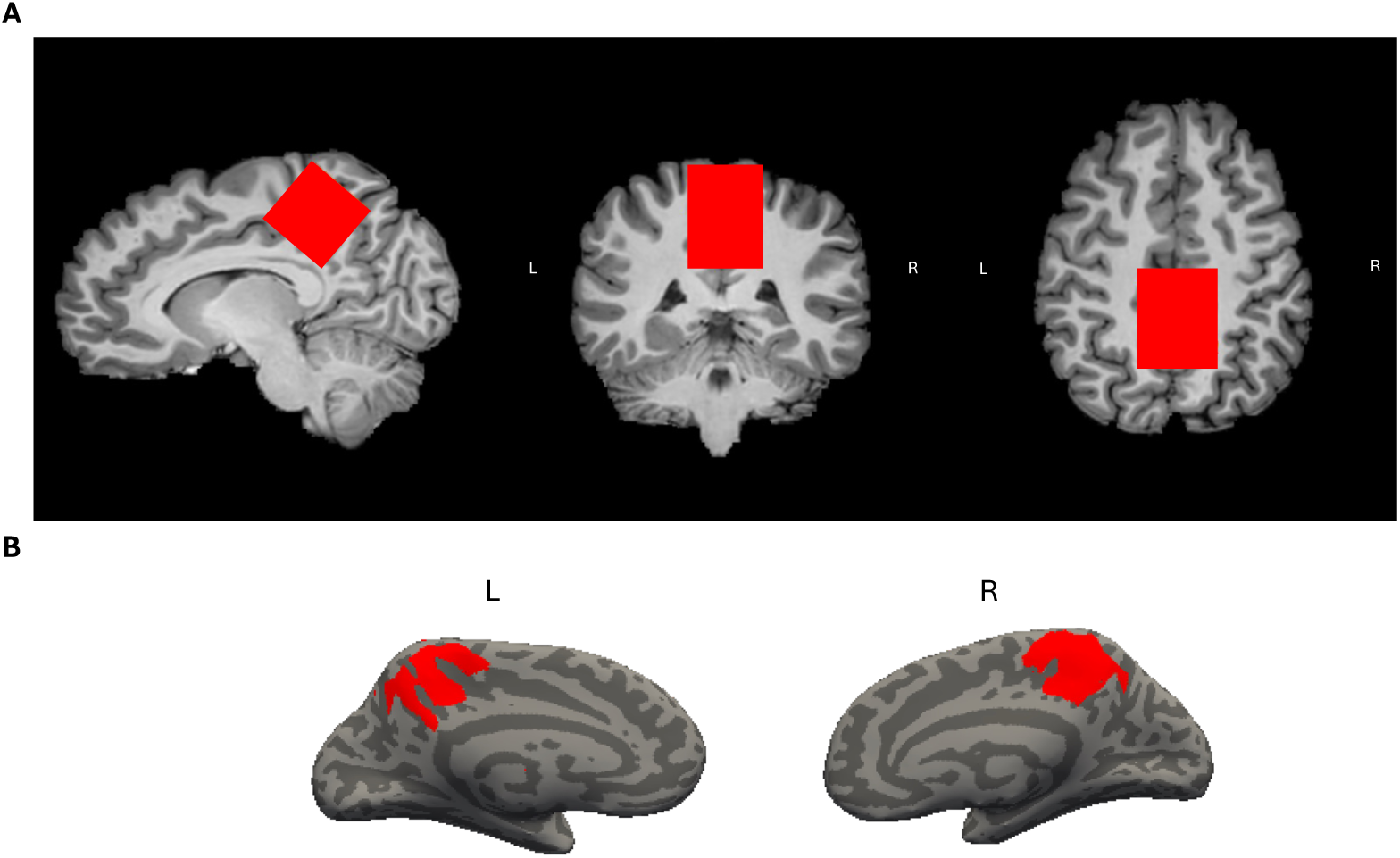
(A) Example PPC voxel mask mapped onto the corresponding T_1_-weighted image. (B) The same PPC voxel mask projected onto an inflated cortical surface of the left and right hemisphere (cortical surfaces were obtained using the FreeSurfer).

### MRS data processing

Raw MRS data were processed using Osprey (Version 2.4.0; (Oeltzschner et al., 2020); version 2022a). First, raw data were eddy current-corrected using the water reference followed by frequency and phase correction using robust spectral registration (Mikkelsen et al., 2020; Near et al., 2011; Oeltzschner et al., 2020), Fourier transformation and water signal removal. Edit-ON and edit-OFF spectra averages were subtracted to resolve GABA+ (at 3.02 ppm) in the difference spectrum (GABA-DIFF). Edit-ON and edit-OFF transients were added to yield the SUM spectra. Average GABA-DIFF and SUM metabolite spectra were modelled using a TE-specific simulated basis set and a flexible spline baseline available on Osprey and based on MRS vendor, pulse duration and scan sequence parameters (Oeltzschner et al., 2020; Simpson et al., 2017). Basis sets for macromolecule and lipid contributions were integrated as Gaussian basis functions (Oeltzschner et al., 2020). Difference and SUM spectra were modelled between 0.5 ppm and 4 ppm with linear baseline correction and a knot spacing of 0.55 ppm according to the Osprey model algorithm (Oeltzschner et al., 2020).

GABA+ was quantified in the difference spectra, while total NAA (tNAA), total Cho (tCho), total Cr (tCr), mI, and Glx were quantified in the SUM spectra. Note GABA signals are quantified as GABA+ due to contamination from overlapping macromolecular signals, while glutamate is commonly reported as Glx (a combined measure of glutamate and its precursor glutamine) due to significant spectral overlap (Harris et al., 2017; Mullins et al., 2014; Puts & Edden, 2012). Average spectra are shown in Figure 2. The Osprey co-registration module (via SPM version 12) was used to register MRS to the T_1_-weighted images acquired at the scan and segment the voxel volume into gray matter fraction (fGM), white matter fraction (fWM) and cerebrospinal fluid fraction (fCSF). Outputs were visually inspected to ensure accurate localization of the MRS voxel. Segmented T_1_-weighted images were used to obtain tissue-composition-corrected water-scaled estimates of metabolite concentrations in institutional units (IU), using three approaches: first, concentrations are scaled according to the assumption that metabolite concentrations in CSF are negligible (Gasparovic et al., 2006; *CSF-corrected metabolite levels*). Second, metabolite and tissue-water T_1_ and T_2_ relaxation corrections were applied (Gasparovic et al., 2006; Oeltzschner et al., 2020; *Tissue-corrected metabolite levels*). Finally, “alpha correction” of GABA+ concentrations was performed, with the assumption that GABA+ concentration is two times more concentrated in GM compared to WM (Harris, Puts, Barker, et al., 2015; Jensen et al., 2005; *alpha-corrected GABA+ levels*). Metabolite concentrations are not reported relative to total creatine (e.g. tCho/tCr; creatine-scaled), as in our previous work we show that creatine levels are unstable across the developmental stage studied (Thomson et al., 2024). Instead, we focus interpretation on estimated *tissue-corrected* metabolite concentrations (IU; and *alpha-corrected* for GABA+). CSF-corrected metabolite associations are also reported in the Supplementary Materials. T_1_-weighted images and then voxel masks were also registered to a standard MN1152 T_1_-weighted 1mm brain anatomical image for the creation of voxel heat plots (Figure 2).

**Figure 2.**
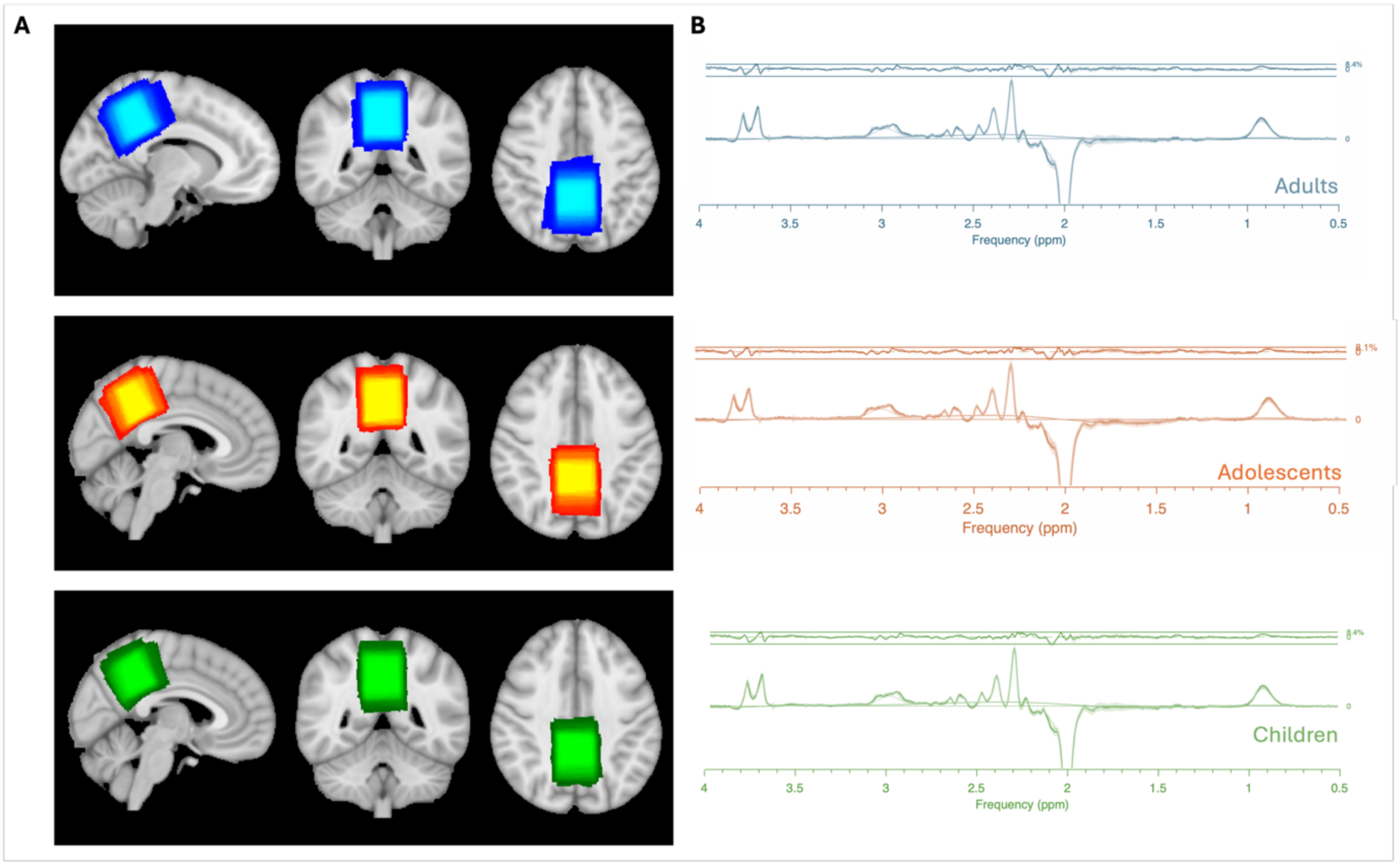
(A) Standard T1-weighted image displaying overlap heat maps of MRS voxels in the PPC across age groups: adults (blue), adolescents (orange), and children (green). Overlap maps were generated by registering individual T1-weighted images and voxel masks to a standard space (MNI152 T1-weighted 1mm brain) and computing voxel overlap. The resulting heat maps are overlaid on the standard anatomical template. (B) Mean MEGA-PRESS GABA-DIFF spectra per age group, showing model fit, baseline, and median residuals (error bars). Ribbon plots indicate the standard deviation of raw spectra fit to basis sets across all spectra.

### Statistical analysis

Analysis was performed in R (Version 4.2.2) using the *lm* function. To observe the effect of metabolite concentrations on structural metrics, linear regression performed with metabolite concentration, sex, age, IQ and estimated total intracranial volume (eTIV) as predictors and the structural metric as the dependent variable (structural metric ∼ metabolite + age + IQ + sex + eTIV). In doing so, we aimed to observe associations between structure and metabolites independent of potential effects of IQ, sex, age and whole brain volume on PPC macrostructure/brain volumes. Controlling for eTIV is common practise for regional brain volume and cortical thickness association studies (see discussion; (Backhausen et al., 2022, 2022; L. Jiang et al., 2016; Voevodskaya et al., 2014). IQ was included as this has previously been associated with cortical thickness variation (Menary et al., 2013). Note that, for surface area metrics, mean cortical surface area was used in place of eTIV in regression models. Partial residual plots showing the effect of metabolite concentration on whole brain/voxel structural metrics (with age, sex, IQ and eTIV held constant) were created using the *effects* package function *effect* (Fox & Weisberg, 2011, 2018).

To correct for the multiple comparisons during regression analysis of metabolite-structural associations (15 metabolites per dependant variable), False Discovery Rate (FDR) correction was applied. FDR controls for the expected proportion of false positives (5% of significant findings). FDR correction also accounts for potential dependencies among statistical tests, a condition that is likely to hold in this case, as PPC metabolites concentrations are metabolically associated. Statistical significance of adjusted p values is reported at two levels: p_adjusted_ < 0.05 and p_adjusted_ < 0.01.

## Results

### Associations between PPC voxel structural metrics and metabolite levels

To assess the consistency with previously characterised age-related changes in cortical macro-structure (Baik et al., 2023; Ducharme et al., 2016; Frangou et al., 2022; Koolschijn & Crone, 2013; Muftuler et al., 2011), we first evaluated whether our PPC voxel macrostructural metrics associated with age. Age significantly and negatively associated with PPC voxel mean cortical thickness, cortical volume, cortical area and LGI when controlling for eTIV (Supplementary Figure 2).

Next, we evaluated associations between PPC voxel structural metrics and metabolite levels, controlling for eTIV, sex, IQ and age. A significant positive linear association was observed between PPC voxel mean cortical thickness and tissue-corrected Glx (beta = 0.019, p_adjusted_ < 0.01; Figure 3). Tissue-corrected Glx also significantly positively associated with voxel cortical volume (beta = 0.019, p_adjusted_ < 0.05; Figure 5) and mean voxel LGI (beta = 0.031, p_adjusted_ < 0.01; Figure 6). Tissue-corrected tCr significantly negatively associated with voxel cortical area (beta = -0.0094, p_adjusted_ < 0.05; Figure 4). Note associations between PPC voxel bulk tissue content (GM, WM and CSF) and metabolite levels are reported in Supplementary Figure 3.

**Figure 3.**
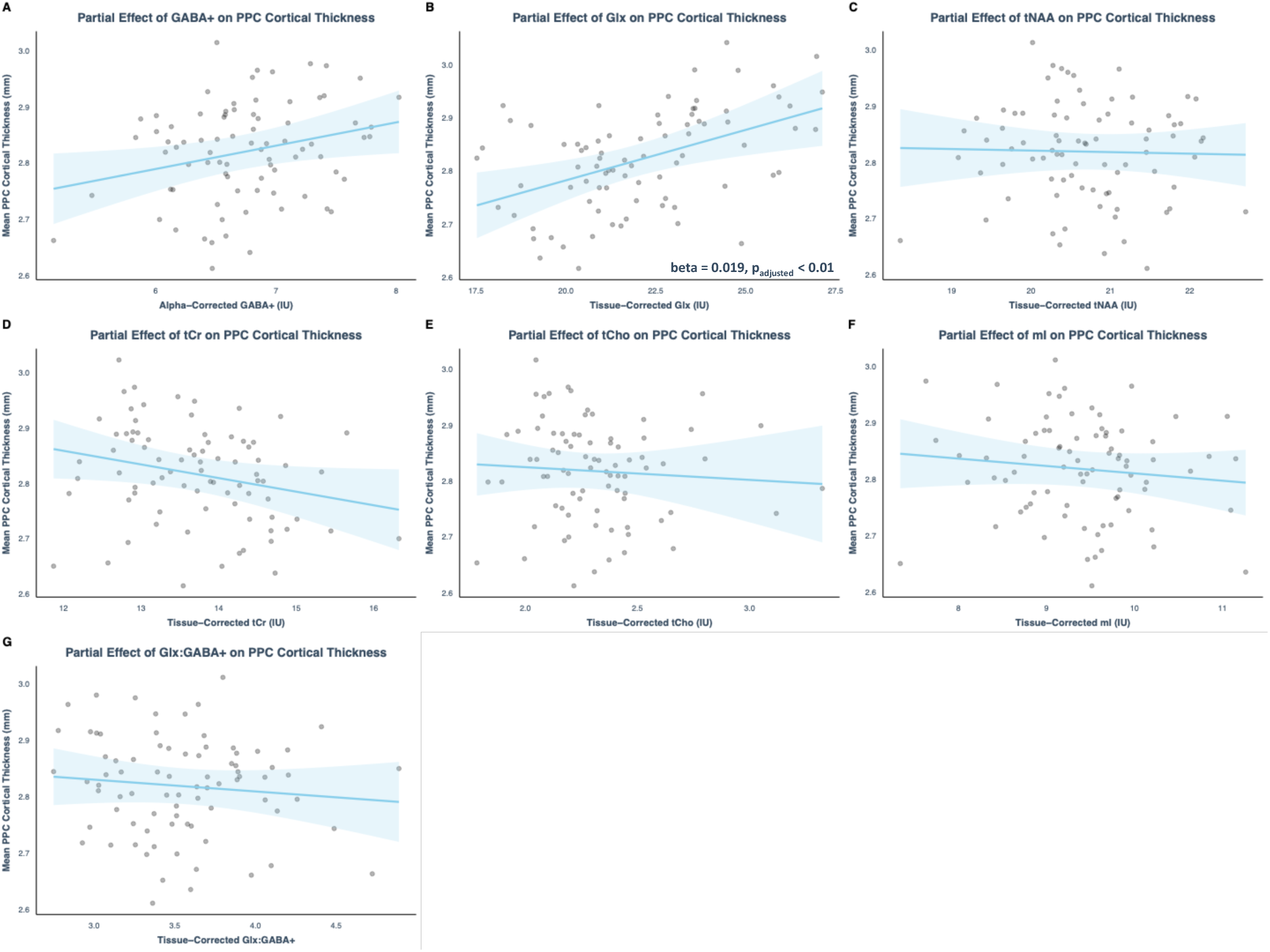
PPC cortical thickness. Partial regression plot shows cortical thickness (mm) predicted by tissue-corrected metabolite concentration (IU) holding participant age, sex, IQ and eTIV constant. Blue shading represents the 95% confidence interval for the partial regression prediction. Points represent individual partial residuals (effect of age, sex, IQ and eTIV regressed out). Significant beta coefficient estimates are shown.

**Figure 4.**
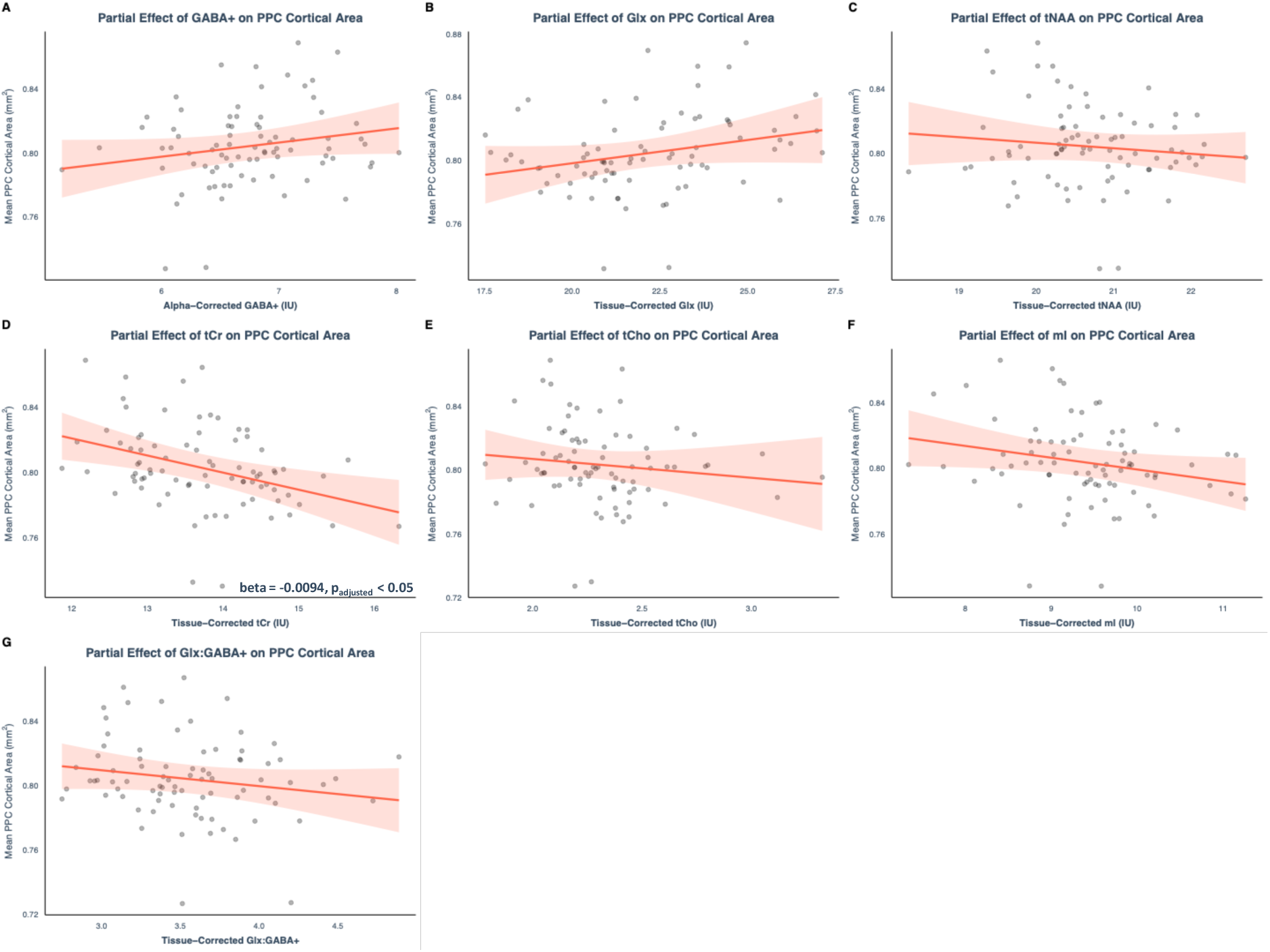
PPC cortical area. Partial regression plot of PPC voxel mean cortical area (mm^2^) predicted by tissue-corrected metabolite concentration (IU) holding participant age, sex, IQ and eTIV constant. Red shading represents the 95% confidence interval for the partial regression prediction. Points represent individual partial residuals. Significant beta coefficient estimates are shown.

**Figure 5.**
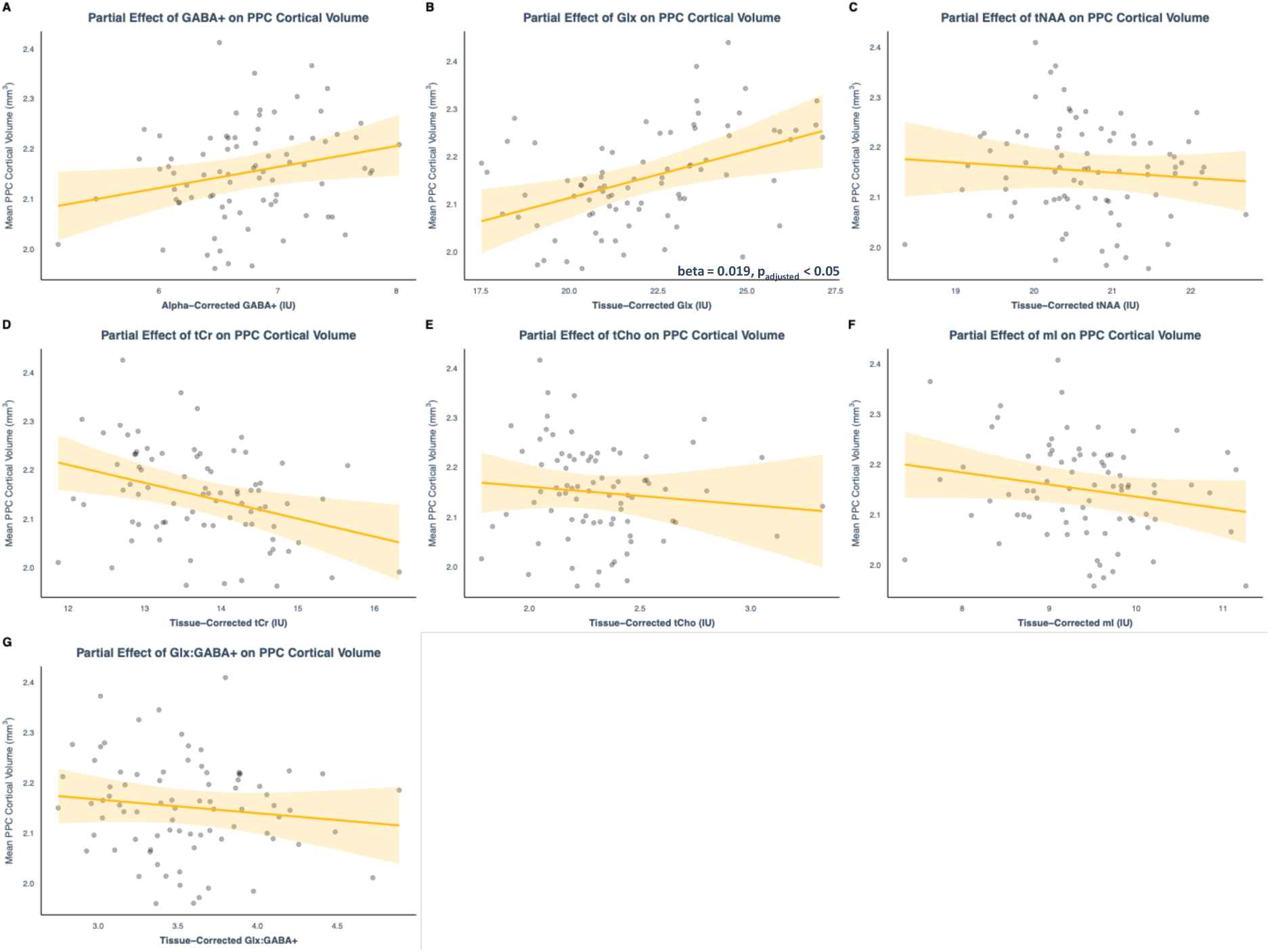
PPC cortical volume. Partial linear regression plot of PPC voxel mean cortical volume (mm^3^) predicted by tissue-corrected metabolite concentration (IU) holding participant age, sex, IQ and eTIV constant. Yellow shading represents the 95% confidence interval for the partial regression prediction. Points represent individual partial residuals. Significant beta coefficient estimates are shown.

**Figure 6.**
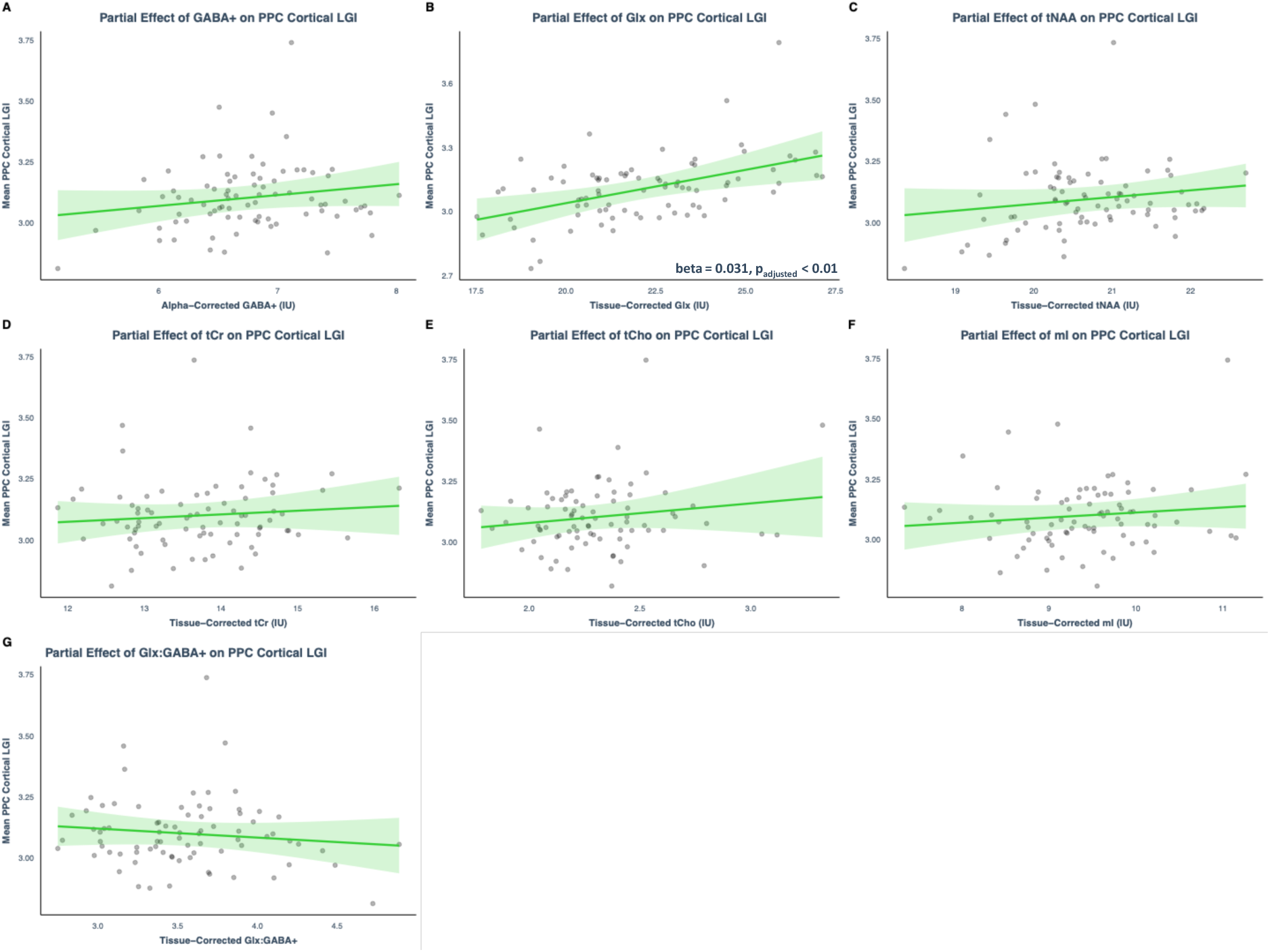
PPC LGI. Partial regression plot of PPC voxel cortical LGI ratio predicted by tissue-corrected metabolite concentration (IU) holding participant age, sex, IQ and eTIV constant. Green shading represents the 95% confidence interval for the partial regression prediction. Points represent individual partial residuals. Significant beta coefficient estimates are shown.

### Associations between whole brain structural metrics and metabolite levels

To determine whether the metabolite-structural associations are region-specific or globally relevant, we also investigate interactions between PPC metabolite concentrations and whole brain structural metrics including white matter volume, subcortical gray matter volume and cortical volume.

We first assessed age-related changes in these metrics. Age significantly negatively associated with global cortical volume and total gray matter volume. Age significantly positively associated with cerebral white matter volume (Supplementary Figure 4).

Next, we evaluated associations between whole brain structural metrics and metabolite levels, controlling for eTIV, sex, IQ and age. tNAA levels significantly positively associated with cerebral white matter volume (beta = 12646.68, p_adjusted_ < 0.05; Figure 7) and cortical volume (beta = 15140.83, p_adjusted_ < 0.01; Figure 9). Cortical volume also significantly positively associated with tissue-corrected Glx (beta = 6547.42, p_adjusted_ < 0.05; Figure 9). Glx:GABA+ ratio significantly negatively associated with subcortical gray matter volume (beta = -2119.67, p_adjusted_ < 0.05; Figure 8).

**Figure 7.**
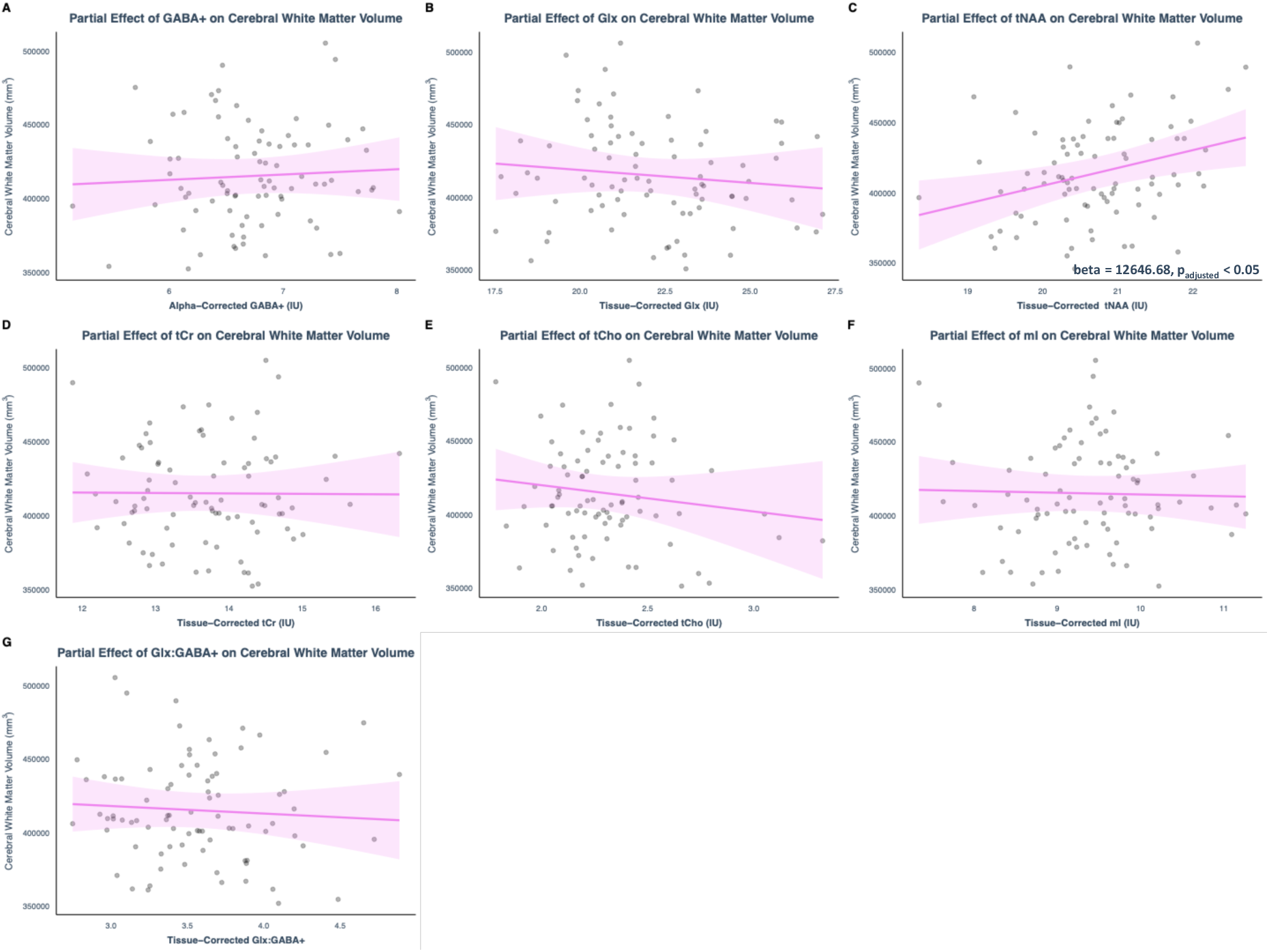
Cerebral white matter volume. Partial linear regression plot of cerebral white matter volume (mm^3^) predicted by tissue-corrected metabolite concentration (IU) holding participant age, sex, IQ and eTIV constant. Pink shading represents the 95% confidence interval for the partial regression prediction. Points represent individual partial residuals. Significant beta coefficient estimates are shown.

**Figure 8.**
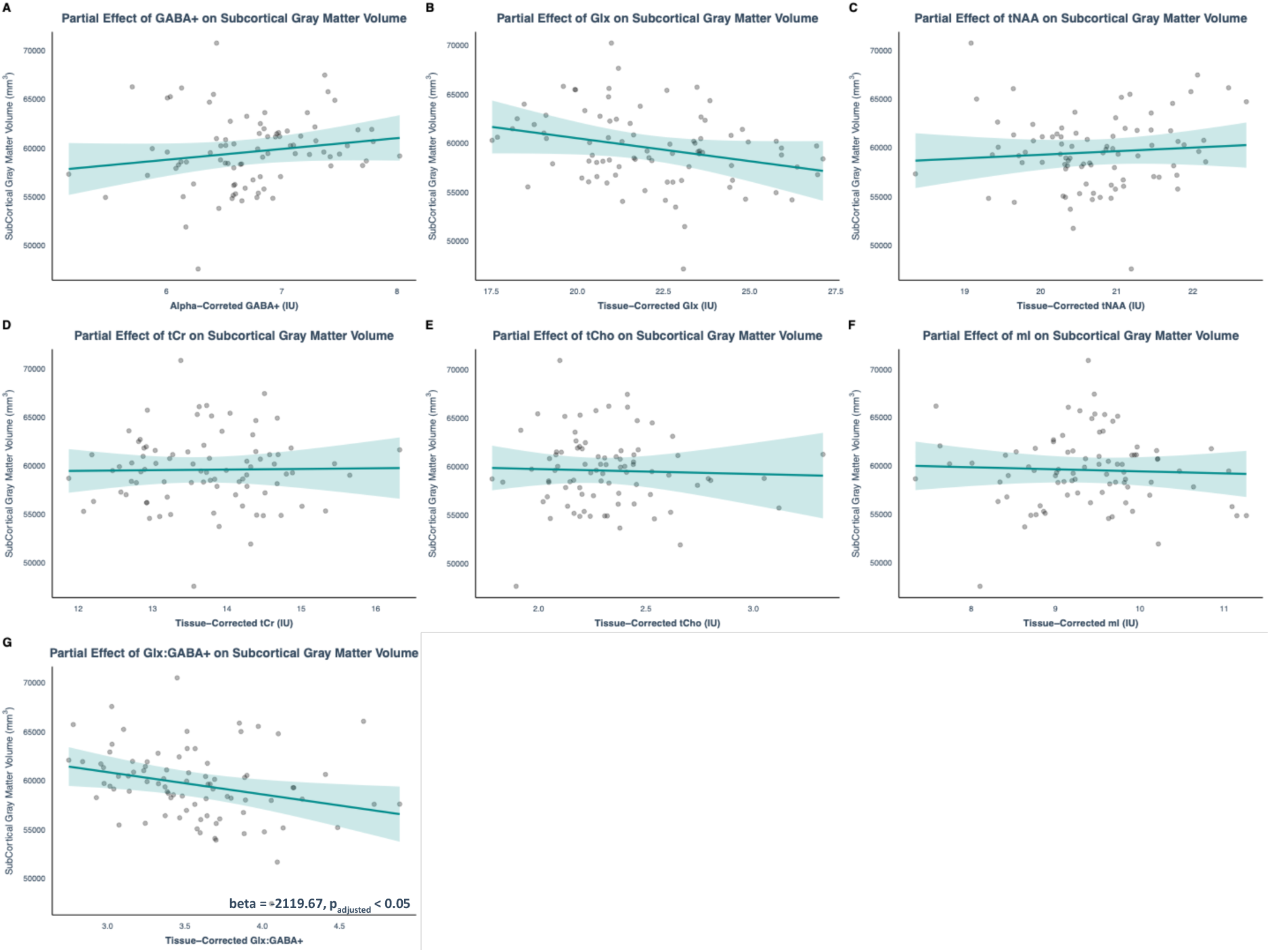
Subcortical gray matter volume. Partial regression plot of subcortical gray matter volume (mm^3^) predicted by tissue-corrected metabolite concentration (IU) holding participant age, sex, IQ and eTIV constant. Blue shading represents the 95% confidence interval for the partial regression prediction. Significant beta coefficient estimates are shown.

**Figure 9.**
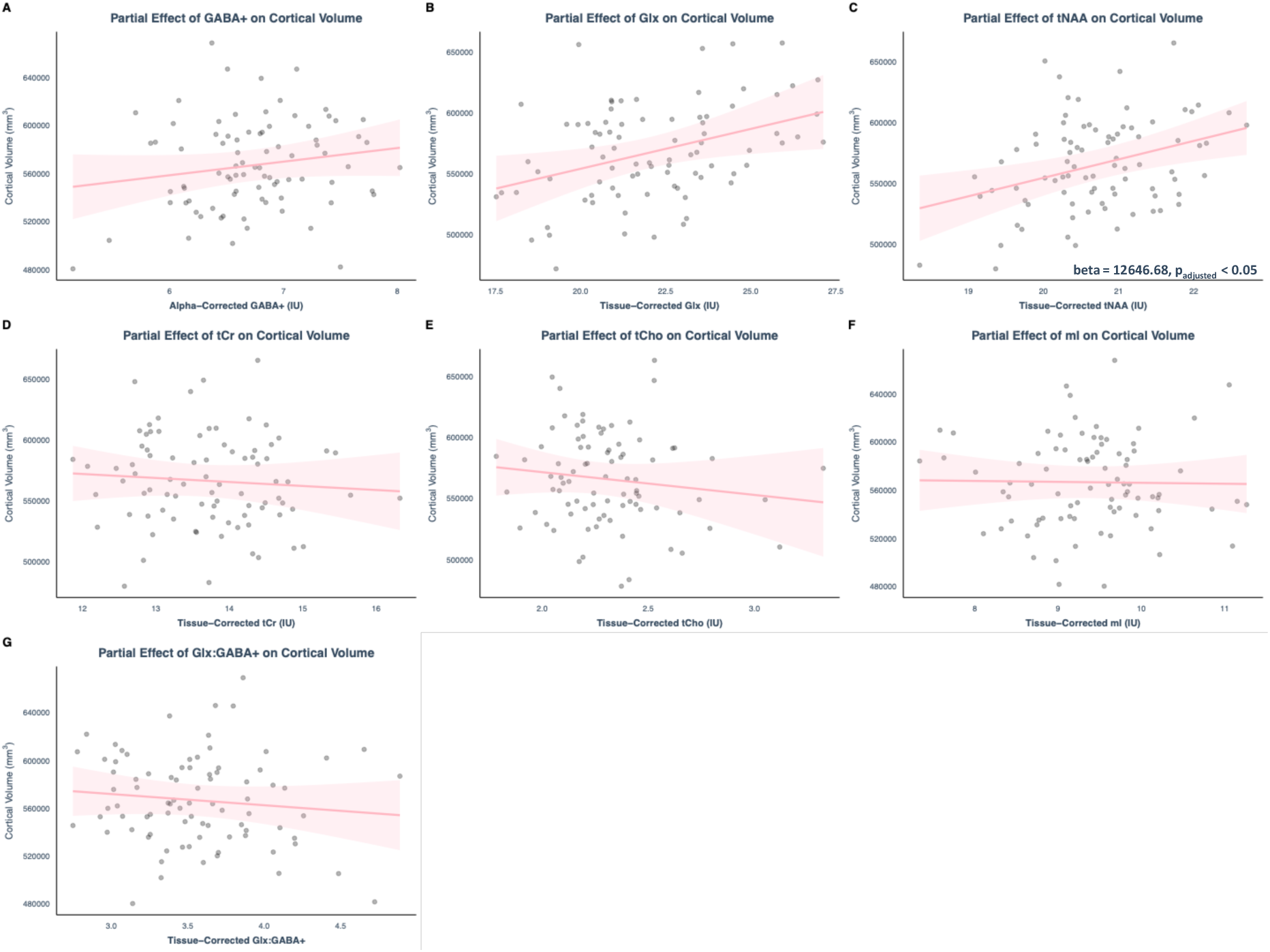
Cortical Volume. Partial linear regression plot of cortical volume (mm^3^) predicted by tissue-corrected metabolite concentration (IU) holding participant age, sex, IQ and eTIV constant. Note GABA+ concentrations were also alpha corrected. Points represent partial residuals. Significant beta coefficient estimates are shown.

## Discussion

In this study, we present the first investigation of the associations between MRS-measured metabolite concentrations and voxel macrostructural metrics in a PPC voxel obtained from a developmental cohort spanning 5 – 35 years of age. Our findings provide novel insights into the neuroanatomical correlates of changes in brain chemistry across childhood to early adulthood, with implications for the interpretation of MRS-measured metabolite levels, and their macrostructural links, in both typical and atypical brain development.

### PPC voxel cortical thickness, area and volume

First we confirmed that cortical thickness, area and volume in the PPC decreases across childhood to early adulthood, consistent with cortical thinning observed in the parietal cortex and multiple other brain regions across the same period (Baik et al., 2023; Ducharme et al., 2016; Frangou et al., 2022; Koolschijn & Crone, 2013; Muftuler et al., 2011). We show that PPC cortical thickness and volume positively associated with Glx concentrations across development, independent of age. Previous studies in adolescence and adulthood report no significant reductions in cortical neuron density (Freeman et al., 2008; la Fougère et al., 2011), despite this, both cortical thickness and Glx decrease across the period studied (Thomson, Pasanta, et al., 2024). Thus, individual differences in neuron density is likely not the sole driver of this association. Instead, these findings likely reflect neuronal and glial cell microarchitectural differences that contribute to individual variability in both cortical macro-structure and Glx across development (Sailasuta et al., 2008; Thomson, Hwa, et al., 2024).

More specifically, histological studies link cortical thinning to changes in neuronal dendritic arbour complexity, which is remodelled by synaptic pruning across the period studied as the brain undergoes circuit refinement (Frangou et al., 2022; Huttenlocher, 1979; Huttenlocher & Dabholkar, 1997; Madre et al., 2020; Petanjek, Judaš, Šimić, et al., 2011). Gene association studies support this, linking genes involved in dendritic spine formation and stability in cortical pyramidal neurons to regional variations in cortical thickness (Parker et al., 2020; Shin et al., 2018; Z. Zhou et al., 2023). Dendritic spines, small protrusions on the dendritic shaft where the majority of excitatory synapses in the cortex are located (Amaral & Pozzo-Miller, 2009), are primary sites of developmental synaptic pruning (Petanjek, Judaš, Šimić, et al., 2011). Given that glutamate, the brain’s primary excitatory neurotransmitter, plays central roles in synaptic activity at dendritic spines, Glx concentrations are likely also related to variations in dendritic arborisation and spine stability. This is supported by findings that Glx concentrations positively correlate with local synaptic density, as estimated using Positron Emission Tomography (PET) to detect radiolabelling of synaptic vesicle glycoproteins (SV2A; (Onwordi et al., 2021). As such, individual differences in dendritic arbour complexity, mediated by synaptic pruning across the developmental period studied, likely contributes to differences in both Glx concentrations and cortical thickness and so the observed association between these traits.

Furthermore, synaptic pruning mechanisms have been linked to glutamatergic signalling (Cachope & Pereda, 2021; Lohmann & Kessels, 2013; Narushima et al., 2016; Surdin et al., 2023). For example, microglia cells engulf synaptic components for pruning, their activity regulated by metabotropic glutamate receptors (Bie et al., 2019; Kim et al., 2017). Given that microglial expressed genes have also been implicated in cortical thickness variation (Parker et al., 2020; Shin et al., 2018; Z. Zhou et al., 2023), it is plausible that Glx concentrations regulate microglial synapse elimination processes on dendritic arbours, thereby driving cortical thickness variation. Further research is required to evidence this and to understand how MRS neurochemical dynamics influence or are influenced by micro and macro-structural brain differences.

Note that cortical thickness differences have also been linked to the myelination of deep intracortical fibres. Cortical thinning during childhood and adolescence is parallel with widespread increases in the fractional anisotropy of major white matter tracts (a structural MRI marker also), indicative of protracted nerve fibre myelination and alignment (Coll et al., 2020; Grydeland et al., 2013; Kinney et al., 1988; Lebel et al., 2008). Myelination may alter MR signal intensity of the deep cortical layers, contributing to the age-related reductions in cortical thickness estimations (B. L. Miller et al., 1996; D. J. Miller et al., 2012; Natu et al., 2019; Paus et al., 2008; Veenstra-VanderWeele et al., 2017; Vidal-Pineiro et al., 2020). Given the positive relationship between Glx concentrations and voxel GM volume, individual differences in deep layer myelination may also contribute to the association between cortical thickness and Glx across the period studied (Zhang & Shen, 2015).

Finally, while the mechanisms underlying cortical area differences are less understood (Norbom et al., 2021), we find a negative association between PPC cortical area and creatine levels. The lack of a Glx-cortical area association, combined with the specific link between creatine and cortical area, suggests that distinct mechanisms govern cortical area and thickness in childhood and adolescence, as supported by previous neuroimaging and genetic studies 28/04/2025 15:03:00. Furthermore, given creatines role in the cellular energy buffering (creatine-phosphocreatine flux; Grasby et al., 2020; Wallimann et al., 1992), mechanisms that drive cortical area variation are suggestively more energetic. Since cortical volume reflects both thickness and area, it is likely shaped by a combination of genetic and environmental factors affecting these traits (Norbom et al., 2021). However, its significant association with Glx points to mechanisms that are more aligned with cortical thickness changes.

### PPC voxel LGI

We find that PPC cortical LGI significantly decreased across the developmental period studied. Changes in LGI were notably more gradual than changes in cortical thickness, consistent with findings in similar developmental cohorts in frontal and temporal regions (Klein et al., 2014; White et al., 2010). The folding or gyrification of the brain predominantly occurs prenatally but continues to adulthood, while the LGI ratio itself becomes relatively stable from the first year of life after birth (White et al., 2010). Gyrification is thought to be essential for the creation of efficient neuronal networks by reducing the distance between neighbouring brain regions to increase communication speeds (White et al., 2010). We find that PPC cortical LGI, similar to cortical thickness, is positively associated with tissue-corrected Glx. Glutamatergic signalling has roles in neuronal migration and axon pathfinding, specifically tuning neuronal responses to axon guidance cues (Kaindl et al., 2008; Luhmann et al., 2015; Tran et al., 2024). Such processes are fundamental to developing early structural connectivity which, in turn, influences cortical folding forces and patterns (Ecker et al., 2016; Nie et al., 2012; Ouyang et al., 2023; Ronan et al., 2014). While mechanisms underlying the subtle changes in LGI observed in our study likely differ from those related to the establishment of gyrification during the prenatal and early post-natal period (Hasan et al., 2011), this association between LGI and Glx might reflect differences in gyrification which were established and associated with glutamatergic mechanisms early on in development and remain detectable in the study period.

Alternatively, changes in LGI during childhood and adolescence have previously been associated with the same mechanisms underpinning changes in cortical thickness. This include dendritic remodelling (White et al., 2010) and intra-cortical fibre myelination, with the diffusivity of white matter fibre tracts specifically associated with individual differences in regional LGI (Ecker et al., 2016). Individual microstructural differences across development may thus alter folding forces, leading to cortical ‘flattening’ observed across the period studied, as well the association between LGI and Glx (Ecker et al., 2016; White et al., 2010).

### Whole brain volumes

PPC biochemistry may interact with broader subcortical and cortical structures. Thus, we chose to also observe interactions between PPC metabolite concentrations and whole brain structural volumes, with significant interactions discussed below.

NAA, the major component of the MRS total NAA signal, is metabolised in oligodendrocytes to support steroid synthesis for the maintenance of myelin sheath integrity (Baslow et al., 1999; Hagenfeldt et al., 1987; Madhavarao et al., 2004; Moffett et al., 2007), a basis for our observations that cerebral white matter volume associated with PPC total NAA levels. However, the absence of a relationship between the white matter fraction *within* the PPC voxel and NAA concentrations, alongside the lack of age-related changes in PPC NAA concentrations despite increases in voxel white matter fraction (Thomson, Hwa, et al., 2024), indicate that local NAA levels may reflect additional biological processes beyond white matter integrity. For example, NAA is highly concentrated within neurons and glial cells, synthesised in the mitochondria where it is utilised in several energetically favourable metabolic pathways (Clark et al., 2006; Patel & Clark, 1979). PCC total NAA may thus also capture individual differences in neuronal and glial mitochondrial distribution, density and metabolic processes, all of which have been implicated in synaptogenesis, synaptic stability, and plasticity across brain development (Markham et al., 2014; Picard & McEwen, 2014; Rangaraju et al., 2019).

Subcortical gray matter volume was negatively associated with PPC Glx:GABA ratio, a proxy measure for E/I balance which decreases across the period studied (Thomson, Hwa, et al., 2024). This association may reflect the shaping of cortical activity by subcortical afferents. Deep cortical layers receive long-range inputs from subcortical thalamic nuclei. Subcortical inputs form stronger and denser excitatory synapses onto interneurons compared to pyramidal cells, leading to feedforward inhibition (Cruikshank et al., 2007; Isaacson & Scanziani, 2011; Li et al., 2014). Within cortical layers, pyramidal cells and interneurons are highly interconnected, forming local inhibitory feedback loops. Combined, feedforward and feedback mechanisms tightly couple cortical excitation (E) and inhibition (I), maintaining E/I balance (Cruikshank et al., 2007; Hirata & Castro-Alamancos, 2010; Isaacson & Scanziani, 2011; Li et al., 2014; Mayer et al., 2019). In mice lacking thalamic-cortical interactions, deep layer cortical neurons display atypical dendritic arbours, reductions in synaptic density and altered topographical organisation (Antón-Bolaños et al., 2018; L. Zhou et al., 2010), thus thalamic-cortical interactions are essential for typical structural development (Antón-Bolaños et al., 2018; Kingsbury et al., 2002). Individual differences in subcortical volume may therefore reflect differences in long-range connectivity from subcortical areas, differentially modulating cortical activity and so underpinning the observed association with the excitatory state of PPC cortical tissue (Glx:GABA+ ratio). This warrants further investigation, particularly given the association between MRS-measured cortical Glx and GABA+ and subcortical-cortical innervation is undefined.

### Limitations

Tissue-corrected metabolite data, which accounts for individual differences in voxel composition (GM, WM, and CSF), is recommended and was reported in this study (Lin et al., 2021). Tissue correction involves applying tissue-specific water and metabolite relaxation corrections (Gasparovic et al., 2006; Lin et al., 2021; Near et al., 2011), but these values are typically derived from adult populations (Oeltzschner et al., 2020). Given that relaxation times likely vary with age, this may introduce bias in metabolite quantification across ages. The accuracy of voxel tissue segmentation and FreeSurfer-derived cortical surface metrics may also be biased by age, with segmentation algorithms primarily trained on adult datasets (Cabezas et al., 2011; Harris et al., 2015). Results were however replicated in metabolite data uncorrected for relaxation values (CSF-corrected only, see Supplementary Materials), partially reinforcing the robustness of the identified associations.

For all analyses, estimated total intracranial volume (eTIV) was included as a covariate to account for co-linear age-related changes in brain volume. Significant associations identified between metabolite concentrations and structural metrics were therefore independent of overall brain volume changes (Backhausen et al., 2022, 2022; L. Jiang et al., 2016; Voevodskaya et al., 2014). Alternative approaches include the use of normalised structural measures (e.g., dividing structural metrics by eTIV), which we chose not to use for several reasons. First, normalisation approaches vary significantly across studies, complicating the interpretation findings. Second, our aim was to investigate the associations between PPC metabolite concentrations and structural metrics. Normalisation might have shifted the focus towards deviations in PPC structure relative to whole-brain averages, obscuring the relationships of interest. Finally, all age-related trends observed in our raw structural metrics (when eTIV is controlled for statistically) are consistent with previous works (Baik et al., 2023; Ducharme et al., 2016; Frangou et al., 2022; Koolschijn & Crone, 2013; Muftuler et al., 2011). However, we acknowledge that alternative approaches may provide complementary perspectives depending on the research focus.

### Conclusion

This study provides novel insights into the relationship between MRS-measured metabolite concentrations and cortical macrostructure across development. We show that PPC cortical thickness, volume and LGI associate with PPC Glx concentrations, likely because differences in cortical microstructures, including dendritic arbour complexity, contributes to differences in cortical macrostructural traits and Glx activity across development. Additionally, PPC metabolite concentrations, particularly the Glx:GABA+ ratio and total NAA, are associated with whole-brain metrics, highlighting the interplay between regional neurochemistry and global structural differences. These findings underscore the importance of accounting for structural variations when interpreting MRS data, especially when comparing metabolite concentrations across groups with (macro)structural differences, such as in neurodevelopmental disorders (Ecker et al., 2022; Hardan et al., 2006; Hyde et al., 2010; You et al., 2024) or across age. Future research should further explore the interdependencies between neurochemical and macrostructural changes to determine whether metabolite alterations are driven by macrostructural changes or if the chemical alterations themselves drive structural changes.

## Acknowledgements

NP and AT are supported through the MRC Centre for Neurodevelopmental Disorders. TA was supported by an MRC Senior Clinical Fellowship [MR/Y009665/1] and the Medical Research Council (MRC) Centre for Neurodevelopmental Disorders [MR/N026063/1]. This work was supported by R01 EB032788, R01 EB016089, R01 EB023963, P41 EB031771 and S10 OD021648, and by the Johns Hopkins Therapeutic Cognitive Neuroscience Fund (Grant Number 80026224) to X.C. and R.L. Engineering Research Council of Canada (XJC; RGPIN-2020-05520) to X.C., Canada First Research Excellence Fund awarded to McGill University for the Healthy Brains for Healthy Lives initiative, and the Canada Research Chairs program.

## Author contribution

**AT:** Data curation, Formal analysis, Methodology, Visualization, Writing – original draft, Writing – review & editing. **DP:** Analysis scripts and Writing—review & editing. **RE & XC**: Conceptualization, Investigation, Resources and Writing—review & editing. **TA & NP**: Conceptualization, Investigation, Writing—review & editing, Supervision, and Resources.

## Conflict of interest disclosure

The authors have no conflicts of interest to disclose.

## Data availability

Data are available through: https://osf.io/uv67s/?view_only=2896964f249f4249bbfdeccb0b96bb73.

Osprey 2.4.0 is available through: https://github.com/schorschinho/osprey

## Notes

### Competing Interest Statement

The authors have declared no competing interest.

