## Supplementary Materials for "Characterising the association between posterior parietal metabolite levels and cortical macrostructure in a cohort spanning childhood to adulthood"

Supplementary Table 1. Results from linear regression of whole brain structural metrics with age, IQ, eTIV and sex. Beta = linear regression predictor coefficient. * Indicates significant p values at; ‘****’ p < 0 ‘***’ p < 0.001 ‘**’ p < 0.01 ‘*’ p < 0.05. R^2^ = adjusted R^2^ obtained from linear modelling.

| **Model** | **Term** | **Estimate** | **P value** | **R squared** | **Significance** |
| --- | --- | --- | --- | --- | --- |
| Cerebral White Matter Volume | Intercept | 83841.62 | 0.046 | 0.56 | * |
|  | age | 2734.44 | 0.00 | 0.56 | **** |
|  | IQ | 332.39 | 0.22 | 0.56 |  |
|  | sex - male | 75.00 | 0.99 | 0.56 |  |
|  | eTIV | 0.17 | 0.00 | 0.56 | **** |
| Subcortical Gray Matter Volume | Intercept | 31266.62 | 0.00 | 0.37 | **** |
|  | age | 2.81 | 0.95 | 0.37 |  |
|  | IQ | 32.98 | 0.27 | 0.37 |  |
|  | sex - male | -117.28 | 0.88 | 0.37 |  |
|  | eTIV | 0.017 | 0.00 | 0.37 | **** |
| Total Gray Matter Volume | Intercept | 395505.50 | 0.00 | 0.66 | **** |
|  | age | -5157.95 | 0.00 | 0.66 | **** |
|  | IQ | 306.91 | 0.34 | 0.66 |  |
|  | sex - male | 10889.41 | 0.21 | 0.66 |  |
|  | eTIV | 0.27 | 0.00 | 0.66 | **** |
| Mean Cortical Thickness | Intercept | 3.41 | 0.00 | 0.72 | **** |
|  | age | -0.021 | 0.00 | 0.72 | **** |
|  | IQ | -0.0016 | 0.036 | 0.72 | * |
|  | sex - male | 0.0037 | 0.86 | 0.72 |  |
|  | eTIV | 5.62E-08 | 0.37 | 0.72 |  |
| Cortex Volume | Intercept | 316316.34 | 0.00 | 0.64 | **** |
|  | age | -4911.05 | 0.00 | 0.64 | **** |
|  | IQ | 141.01 | 0.63 | 0.64 |  |
|  | sex - male | 9821.53 | 0.21 | 0.64 |  |
|  | eTIV | 0.22 | 0.00 | 0.64 | **** |

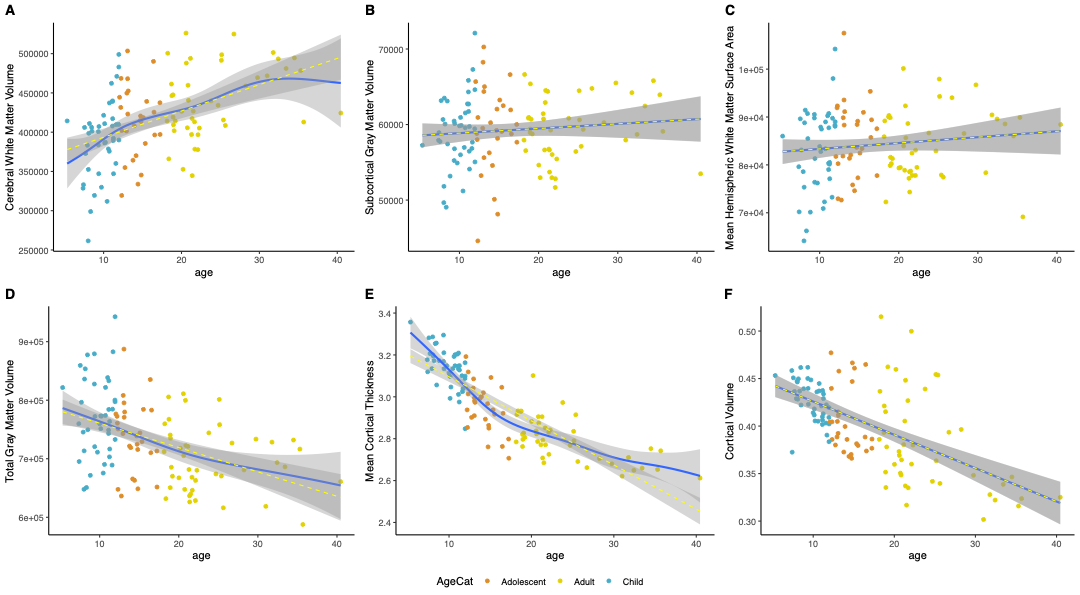

Supplementary Figure 1. Whole brain structural metrics were plotted against participant age. (A) Cerebral white matter volume (B) Subcortical gray matter volume (C) Mean hemispheric white matter surface area (D) Total gray matter volume (E) Mean whole brain cortical thickness (F) Cortical volume. Linear regression (yellow) and non-linear GAM (blue) model fits are shown. Blue point = child, orange point = adolescent, yellow point = adult. Gray bars represent the 95% confidence interval of model fit.

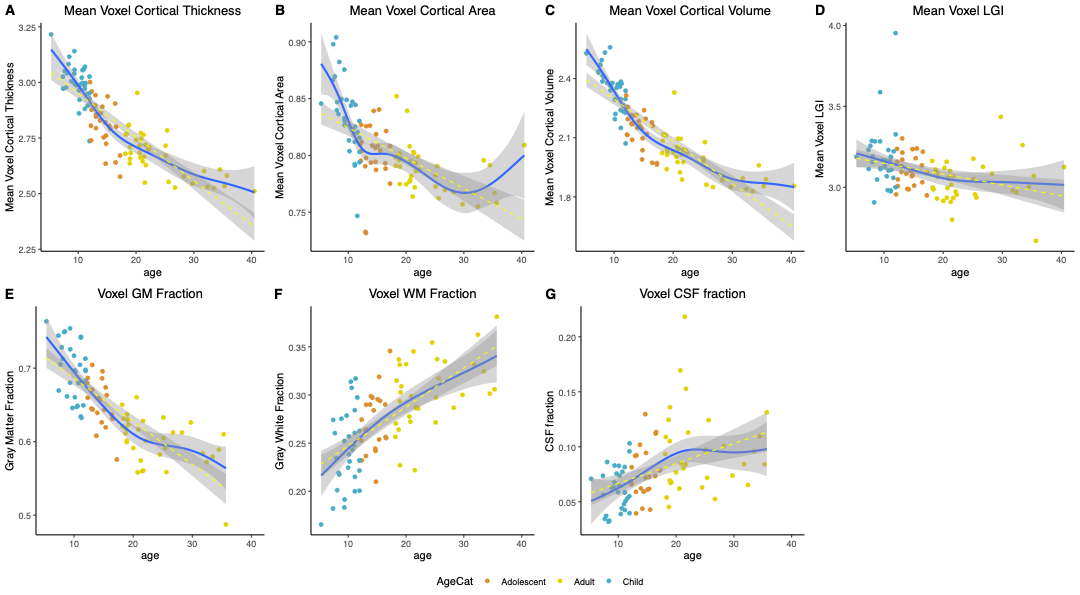

Supplementary Figure 2. PPC voxel cortical surface-based metrics plotted against participant age. (A) Mean PPC voxel cortical thickness (B) Mean PPC voxel cortical area (C) Mean PPC voxel cortical volume (D) Mean voxel local gyrification index (LGI) (E) Voxel GM fraction (F) Voxel WM fraction (G) Voxel CSF fraction. Linear (yellow-hashed line) and non-linear GAM models (blue line) are shown. Blue point = child, orange point = adolescent, yellow point = adult.

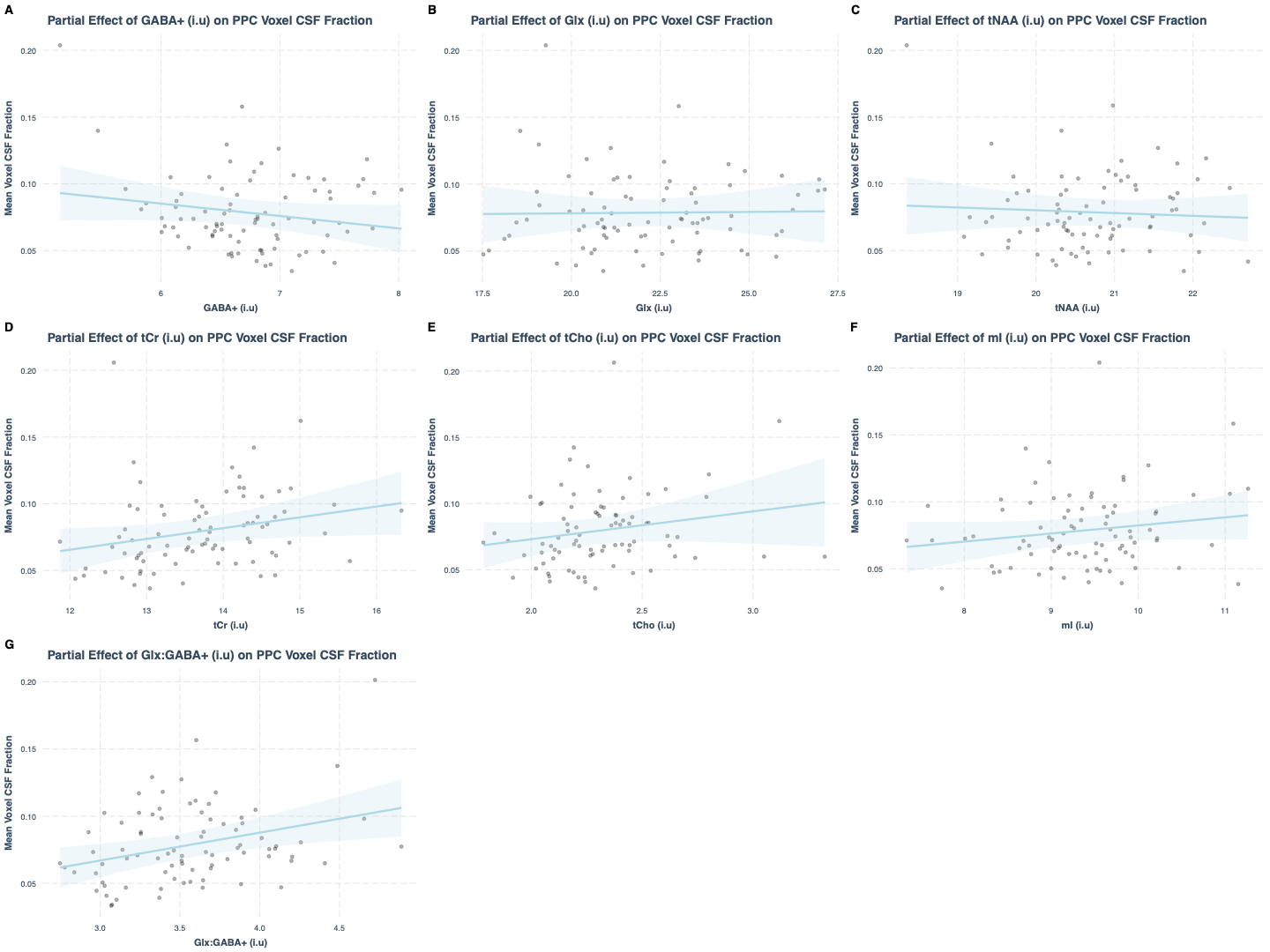

Supplementary Figure 3. Partial linear regression plot of PPC voxel CSF fraction predicted by tissue-corrected metabolite concentration holding participant age, sex, IQ and eTIV constant. Blue shading represents the 95% confidence interval for the partial regression prediction. Points represent individual partial residuals. A significant positive association between voxel CSF fraction and tissue-corrected Glx:GABA+ (beta = 0.021, padjusted < 0.05) was observed. No significant associations between metabolite levels and voxel GM or WM fraction were observed.

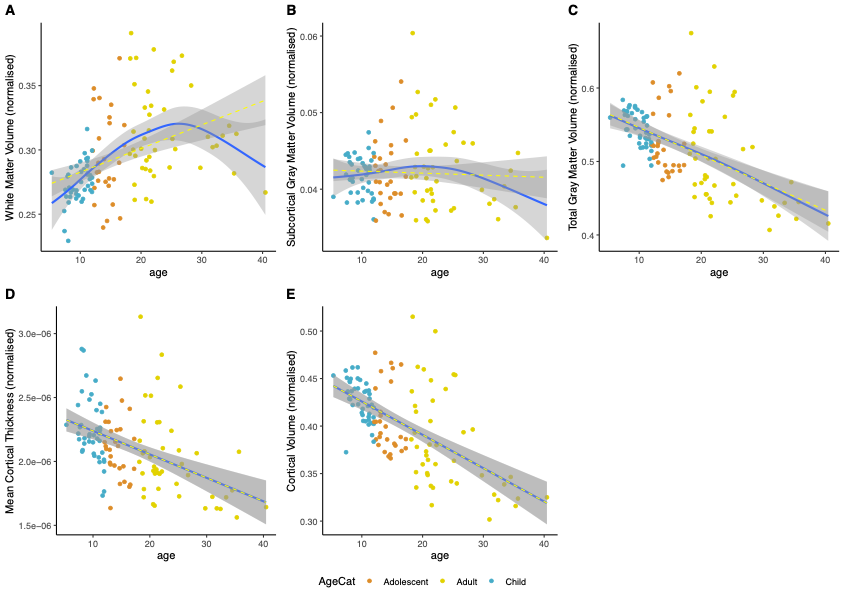

Supplementary Figure 4.Whole brain structural metrics (NORMALISED) extracted were plotted against participant age. (A) cerebral white matter volume (V) subcortical gray matter volume (C) total gray matter volume (D) mean cortical thickness and (E) cortical volume plotted against years of age. Linear (yellow) and non-linear model fits are shown. Blue point = child, orange point = adolescent, yellow point = adult.

CSF-corrected metabolites

A significant positive linear association was observed between PPC voxel mean cortical thickness and CSF-corrected GABA+ (beta = 0.06, p_adjusted_ < 0.05) and CSF-corrected Glx (beta = 0.023, p_adjusted_ < 0.05). CSF-corrected Glx also significantly positively associated with voxel cortical volume (beta_CSF-corrected_ = 0.023, p_adjusted_ < 0.01) and mean voxel LGI (beta_CSF-corrected_ = 0.036, p_adjusted_ < 0.01). CSF-corrected tCr significantly negatively associated with voxel cortical area (beta_CSF-corrected_ = -0.0010, p_adjusted_ < 0.05; Figure 4).

tNAA levels significantly positively associated with cerebral white matter volume (beta_CSF-corrected_ = 13798.82, p_adjusted_ < 0.05) and cortical volume (beta_CSF-corrected_ = 15192.00, p_adjusted_ < 0.01). Cortical volume also significantly positively associated with CSF-corrected Glx (beta = 7832.65, p_adjusted_ < 0.01). Mean global cortical thickness significantly positively associated with CSF-corrected GABA+ (beta = 0.067, p_adjusted_ < 0.05) and CSF-corrected Glx (beta = 0.024, p_adjusted_ < 0.01).
